# Brain-state modeling using electroencephalography: Application to adaptive closed-loop neuromodulation for epilepsy

**DOI:** 10.1101/2024.06.01.596884

**Authors:** Graham W. Johnson, Derek J. Doss, Ghassan S Makhoul, Leon Y. Cai, Camden E Bibro, Danika L. Paulo, Shilpa B. Reddy, Robert P. Naftel, Kevin F. Haas, Mark T. Wallace, Benoit M. Dawant, Angela N. Crudele, Victoria L. Morgan, Christos Constantinidis, Shawniqua Williams Roberson, Sarah K. Bick, Dario J. Englot

## Abstract

The progress of developing an effective closed-loop neuromodulation system for many neurological pathologies is hindered by the difficulties in accurately capturing a useful representation of a brain’s instantaneous functional state. Existing approaches rely on expert labeling of electroencephalography data to develop biomarkers of neurophysiological pathology. These techniques do not capture the highly complex functional states of the brain that are presumed to exist between labeled states or allow for the likely possibility of variation among identically labeled states. Thus, we propose BrainState, a self-supervised technique to model an arbitrarily complex instantaneous functional state of a brain using neural multivariate timeseries data. Application of BrainState to intracranial electroencephalography data from patients with epilepsy was able to capture diverse pre-seizure states and quantify nuanced effects of neuromodulation. We anticipate that BrainState will enable the development of sophisticated closed-loop neuromodulation systems for a diverse array of neurological pathologies.

## INTRODUCTION

Neuromodulation for drug-resistant epilepsy (DRE), and other neurological pathologies, has advanced considerably in the last 15 years.^1^ This includes the ability to monitor the brain for potential biomarkers of active disease states and stimulate based on a feedback loop, i.e. closed-loop neuromodulation.^2,3^ For DRE, closed-loop neuromodulation currently relies on detecting immediate pre-ictal signatures and delivering high-energy stimulation in an attempt to abort seizure progression, coined Responsive Neurostimulation (RNS).^4^ This paradigm has shown efficacy in treating DRE,^5^ and served as the vanguard for the development of closed-loop neuromodulation. Recent insight into the potential mechanism of RNS indicates that interictal stimulation may actually be driving therapy,^6^ and that low-energy stimulation may be the most effective paradigm.^7^ Furthermore, these hypotheses are in alignment with the clinical observation that seizures tend to occur on long-range periodic timescales.^8^ Thus, it is reasonable to hypothesize that a long timescale biomarker that models seizure propensity in a continuous distribution and allows for precision low-energy interictal stimulation may be most effective for device feedback.^9^ Additionally, an ideal biomarker would have the ability to be continuously and automatically updated based on new data (i.e. *adaptable* closed-loop neuromodulation). The development of such a biomarker using intracranial electroencephalography (iEEG) is the scope of this work.

A continuous seizure propensity biomarker is distinct from seizure forecasting.^10,11^ Seizure forecasting provides a unidimensional quantification of seizure likelihood and assumes that the functional state of the brain (i.e. “brain-state”) is always on a linear progression toward a seizure. It is more reasonable to assume that a person with epilepsy vacillates in and out of *different* high-seizure propensity states. However, if a given brain-state does not immediately progress to a seizure, it cannot be labeled by an epileptologist due to lack of a clear electroclinical sign. Seizure forecasting relies on these ictal and preictal labels to build its prediction modeling. In contrast – akin to the concept of a ‘tornado *watch’* describing high-risk conditions for tornadic activity over the timescale of hours versus a ‘tornado *warning’* indicating an actual funnel cloud forming with immanent progression to a tornado – a closed-loop biomarker must allow for the mapping of high-risk brain-states that are presumed to exist over the scale of hours to days (tornado watch) without immediate transformation to seizure activity (tornado warning / funnel cloud). Thus, a more generalized framework, similar to brain-machine interfaces,^12^ that captures seizure propensity in a long timescale brain-state space may be most effective for closed-loop neuromodulation. Importantly, this paradigm must not rely on any labeling to generate its understanding of the brain-state space. To contextualize this framework for DRE, we have formalized a few guiding postulates and their implications to biomarker generation (**Table 1**).

**Table 1:** Epilepsy brain-state postulates.

| Epilepsy Brain-State Postulates |  |
| --- | --- |
| **DEFINITIONS** |  |
| **Latent space:** | The regularized multidimensional space of possible brain-states constructed from previously observed phenomena (e.g. intracranial electroencephalography). |
| **Brain-state/Embedding:** | The exact mapped location within the latent space. Brain-state is synonymous with ‘embedding’ in this context. |
| **Latent distance:** | The mathematical distance between two brain-states in the high-dimensional latent space, as defined by metrics like Euclidean, Manhattan, or angular distance. The higher the latent distance, the more ‘different’ the brain states are. Words like ‘far’ and ‘furthest’ are used to indicate brain-states separated by a large latent distance. |
| **POSTULATES** |  |
| **Postulate 1:** | NO TWO PRE-ICTAL BRAIN-STATES ARE IDENTICAL. |
| **Implication:** | *Exact latent space embedding of previous events only serves as approximate landmark for future events. Furthermore, different types of ictal events (e.g. subclinical, focal aware, focal impaired-aware, focal to bilateral tonic-clonic) likely embed into different latent regions.* |
| **Postulate 2:** | TEMPORAL EVOLUTION FROM PRE-ICTAL TO ICTAL BRAIN-STATE CAN VARY. |
| **Implication:** | *The exact time course of brain-states in latent space before a given seizure cannot be reliably used as a label for latent space construction or post-hoc interpretation. Specifically, not all brain-states adjacent to a previously verified pre-ictal state will evolve into a seizure in the same manner – there is an inherent level of stochasticity that is difficult to model. Thus, latent space construction benefits from a purely unsupervised/self-supervised model to avoid bias.* |
| **Postulate 3:** | THE BRAIN-STATE ‘FURTHEST’ FROM OBSERVED PRE-ICTAL AND ICTAL ACTIVITY IS ILL-DEFINED. |
| **Implication:** | *In a patient currently experiencing multiple seizures, it is likely that few to no ‘stable’ brain-states far from ictal activity are observed and mapped into the latent space. Thus, a theoretical optimal brain-state target for closed-loop neuromodulation is likely not readily defined by previously observable brain states. Furthermore, there is likely a constellation of potentially stable brain states far from potential ictal activity.* |

Artificial intelligence (AI) has been proposed as a solution to biomarker generation, but the concept of how to effectively model long-term brain-states with AI has remained elusive.^13-15^ Furthermore, the optimal signal processing required to convert raw multi-channel one-dimensional iEEG signals into data ready for AI model generation is not well characterized due to presence of high-voltage interictal spiking and ictal activity creating skewed data distributions. Thus, we propose a modeling process that utilizes novel signal processing and normalization, a custom asymmetric variational autoencoder, and a novel loss paradigm to elucidate a generalized model of brain-state that can be used in adaptable closed-loop neuromodulation. With this brain-state biomarker (**Figure 1: Steps 1-3**), a second model within the closed-loop device ecosystem is then trained to deliver the proper low-energy neurostimulation over long time periods to maintain a satisfactory brain-state (**Figure 1: Step 4**). Finally, to adapt to the potential very long-term (weeks) shift in brain-states, the brain-state model can be continuously adapted in the context of new data and observed stimulation efficacy (**Figure 1: Steps 5-6**).

**Figure 1:**
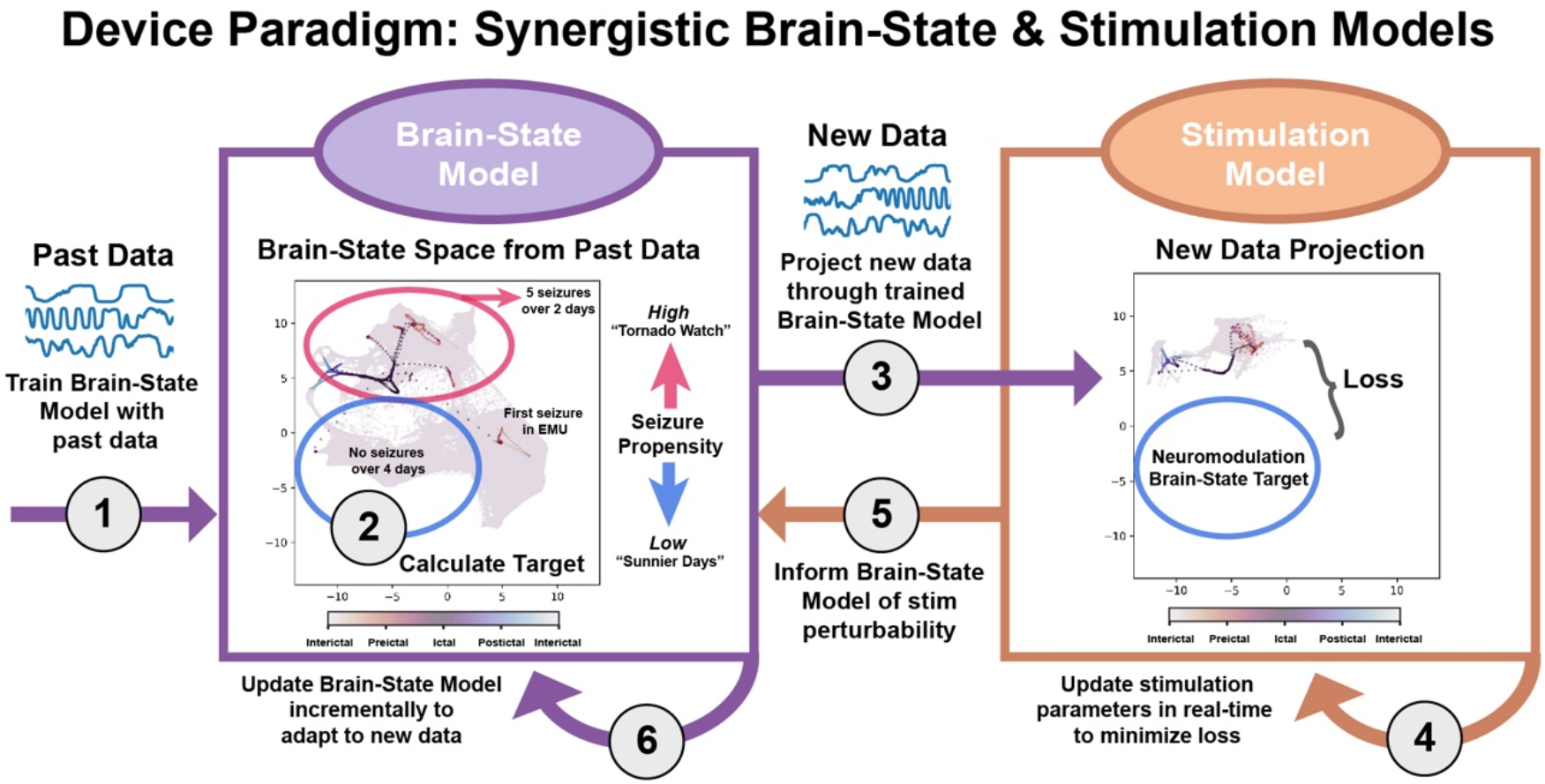
Synergistic adaptable closed-loop neuromodulation models to elucidate brain-state and deliver brain-state modulating low-energy stimulation.

In summary, the future of closed-loop adaptive neuromodulation for drug-resistant epilepsy, and other neurological pathologies, relies on a biomarker that can effectively quantify long timescale disease propensity in a smooth and continuous distribution for effective device feedback. The techniques required to model large-timescale brain-states need not be specific to epilepsy and could be applicable to a broad array of neurological pathologies and neuromodulation conditions. Thus, the techniques described herein will be mostly kept agnostic to disease and focus on the generalized concept of brain-state modeling with multi-channel neurophysiological timeseries. However, we will demonstrate and validate these methods using stereotactic-electroencephalography (SEEG, a subset of iEEG) timeseries from patients undergoing pre-surgical workup for DRE.

## METHODS

As stated, the goal of this work was to develop a generalizable framework of creating a functional brain-state map from high-quality intracranial electrophysiological timeseries for the purpose of serving as a biomarker for closed-loop neuromodulation. To develop and validate the brain-state model architecture, we utilized our cohort of approximately 17,000 hours (16.3 TB of 32-bit precision) of continuous SEEG data from 118 patients with DRE undergoing SEEG presurgical evaluation at Vanderbilt University Medical Center (VUMC). This study was approved by Vanderbilt’s Institutional Review Board and all patients provided informed consent.

### Brain-state modeling

#### Data acquisition

We collected SEEG data using Natus Neuroworks (Middleton, WI, USA) with a sampling rate of 512-2048 Hz. All data was resampled down to 512 Hz then filtered using MATLAB’s (MathWorks inc., Natick, MA, USA) ‘filtfilt’ function to implement a zero-phase shift 5^th^ order Butterworth filter with passbands of 1-59, 61-119, and 121-179 Hz.^16^ An adjacent bipolar montage was applied across all SEEG leads and data was saved as 32-bit floating point precision Python pickle files (**Figure 2A**).

**Figure 2:**
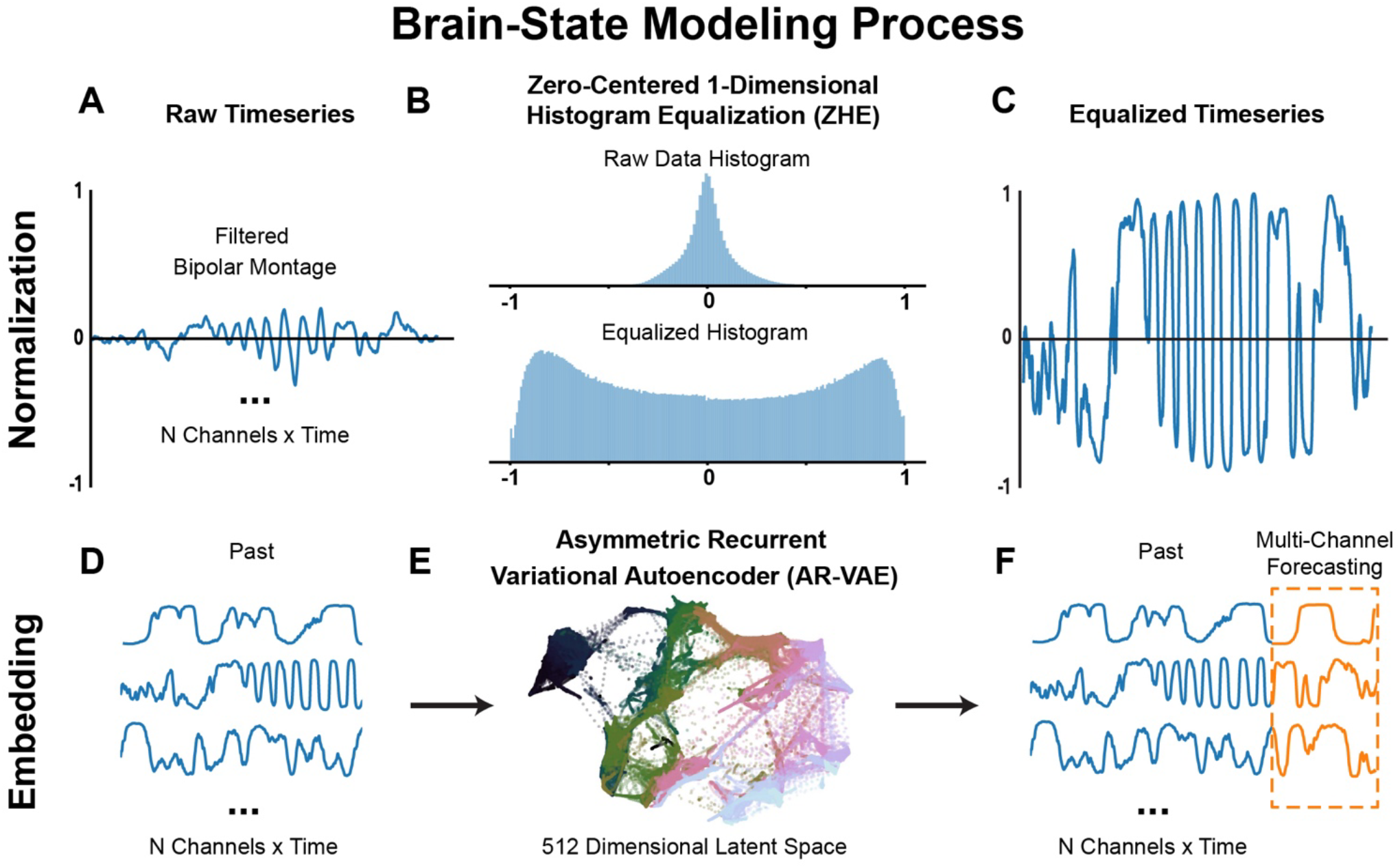
Proposed brain-state modeling process.

#### Zero-Centered One-Dimensional Histogram Equalization (ZHE)

Intracranial electrophysiological timeseries are often low amplitude with rare high amplitude signatures. This can make it difficult for a model to learn nuanced long-range characteristics because the histogram of the data distribution can have long tails (**Figure 2B** top panel) representing rare, but physiologically meaningful high voltage signatures. To provide more evenly distributed data suitable for machine learning models, we implemented a custom Zero-centered one-dimensional Histogram Equalization (ZHE) scheme (**Figure 2B**). This scheme was motivated by traditional histogram equalization for images prior to model training. First, the signal is split into the positive and negative domains, then a 10,000-bin histogram is filled for each domain from 0 to absolute value of highest voltage. Then the linear transfer function from the raw signal to the equalized signal is calculated using the cumulative distribution function for the positive and negative domain separately. The result is a signal that preserves the physiological meaning of zero, but has data evenly distributed between -1 and 1 (**Figure 2C**). A prospective normalization scheme must be applied to not violate causality while validating the model on subsequent data epochs, thus all subject’s timeseries were normalized to the first 24 hours of data. The histograms for subsequent data epochs will, as expected, not be exactly uniform (e.g. **Figure 2B** bottom panel).

#### Brain-state embedding model

The ZHE data are then used to elucidate brain-states by compressing short (0.5 seconds) data epochs (**Figure 2D**) into a 512-dimensional latent space (**Figure 2EB)** that is then used to forecast the immediate future (0.125 seconds) of all channel data simultaneously (**Figure 2F**). This embedding is conducted using a custom Asymmetric Recurrent β-Variational Autoencoder (AR-βVAE), implemented in PyTorch,^17^ and trained on the first 70% of a subject’s data (**Figure 3**). A model was trained for each subject separately to accommodate unique SEEG implants and presumed unique neurophysiological brain-states experienced by each individual.

**Figure 3:**
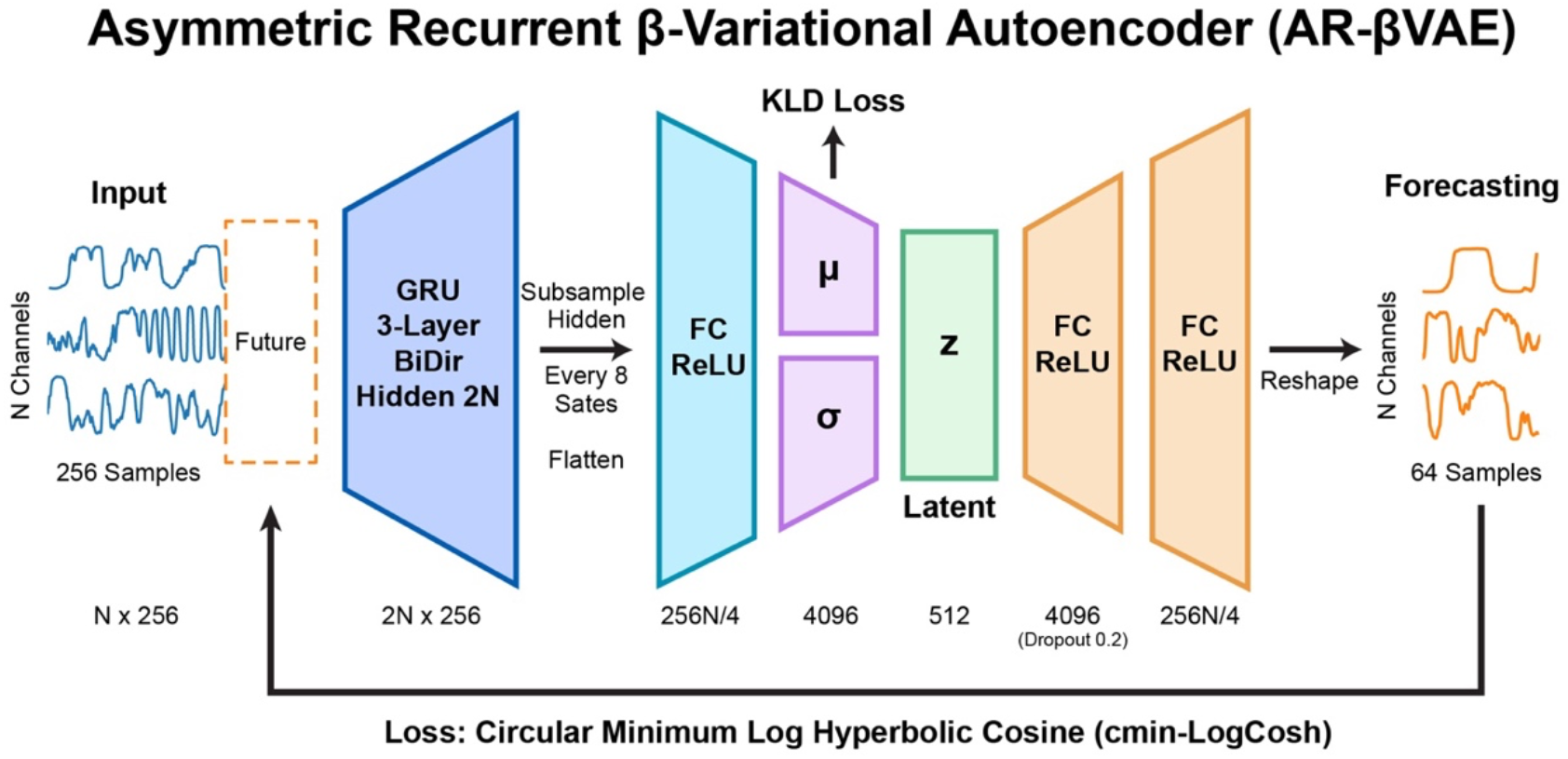
Custom Asymmetric Recurrent Beta-Variational Autoencoder architecture.

The AR-βVAE was architected to accept an N-channel (for our dataset, N ranged from 88-168 channels) by 256 sample (i.e. at 512 Hz sampling rate, 0.5 seconds of multichannel data). The top layer of the model utilizes a Gated Recurrent Unit (GRU) to interface directly with the input data and learn short- and long-range signal features. The GRU is three layers and bidirectional, with a resulting hidden dimension of 2N. The hidden state is then subsampled every eight states. In theory, this helps the GRU to maintain learning capacity and not be forced to forget data motifs across the entire 256 forward and 256 reversed sequence lengths. The subsampled hidden states are then flattened and fed into a β-VAE consisting of a fully-connected layer feeding into mean (μ) and log-variance (σ) layers followed by the standard noise-injection reparameterization trick to get ‘z’, the latent space.^18^ The latent space was regularized during training with Kullback-Leibler (KL) Divergence. The latent space of the variational autoencoder is set to 512 dimensions. The decoder outputs an Nx64 sample forecast on all channels simultaneously. Thus, all Nx256 input timepoints are compressed into 1×512 latent dimensions that best embed the necessary information to forecast the next Nx64 timepoints in the original input signal. A dropout ratio of 0.2 was used on the middle fully connected layer of the decoder to promote concise and meaningful latent embeddings.^19^

#### Circular minimum log hyperbolic cosine loss

Of significance, the decoding portion of the β-VAE is asymmetric because the input GRU layer in the encoder cannot be easily reversed for the decoder. This creates an undesirable situation where slightly shifted input signals could require dramatically different embeddings due to the rigidity of the output fully-connected decoding layers – i.e. small time shifts in the input data could require very different outputs from the final fully connected layers. To accommodate this asymmetric intractability, a custom loss function (**Figure 4**) was developed that allows for a varying time shift of the forecasted signals, termed Circular Minimum Hyperbolic Cosine Loss (cMin-LogCosh). This loss function was based on the hyperbolic cosine loss function as implemented in PyTorch by ‘auraloss’ package.^20,21^ Specifically, the Nx64 predicted values are wrapped in a circle using ‘torch.roll’ in PyTorch with a stride of one sample (**Figure 4A-B**) and the LogCosh loss calculated every stride resulting in 64 individual loss values. The minimum LogCosh loss is deemed the cMin-LogCosh (**Figure 4C**) and returned for backpropagation through the model.

**Figure 4:**
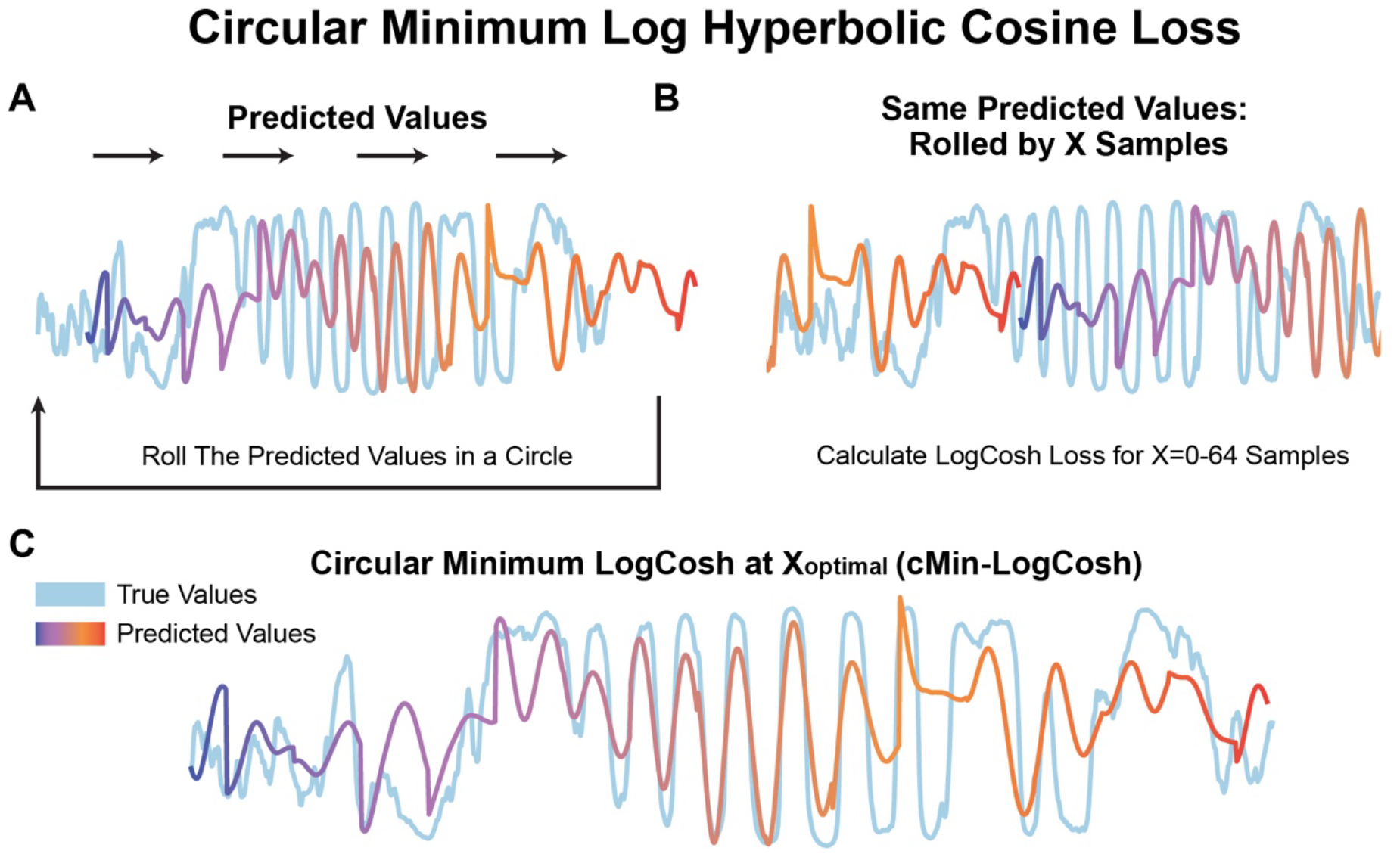
Time-shift invariant loss function for multi-channel timeseries forecasting.

#### Training considerations & Kullback-Leibler / learning rate annealing

Each subject’s model was trained by feeding in random Nx256 epochs of data with a batch size of 64. Each epoch consisted of a total of 640,000 random epochs and the models were each run to a total of 500 epochs, resulting in a total of 320M random epochs used for training. The first 70% of the patient’s data was used for training and the remaining 30% was completely left out and used in the validation sections as described below. In an attempt to maximize biologically meaningful features extracted from the dataset, a sigmoidal KL annealing schedule was implemented with a 40 epoch period (**Figure 5A**).^22^ Additionally, to increase model training stability, a learning rate (LR) annealing with a single epoch saw-tooth waveform and a gamma of 0.1 at 250 of the total 500 training epochs was implemented (**Figure 5B**).

**Figure 5:**
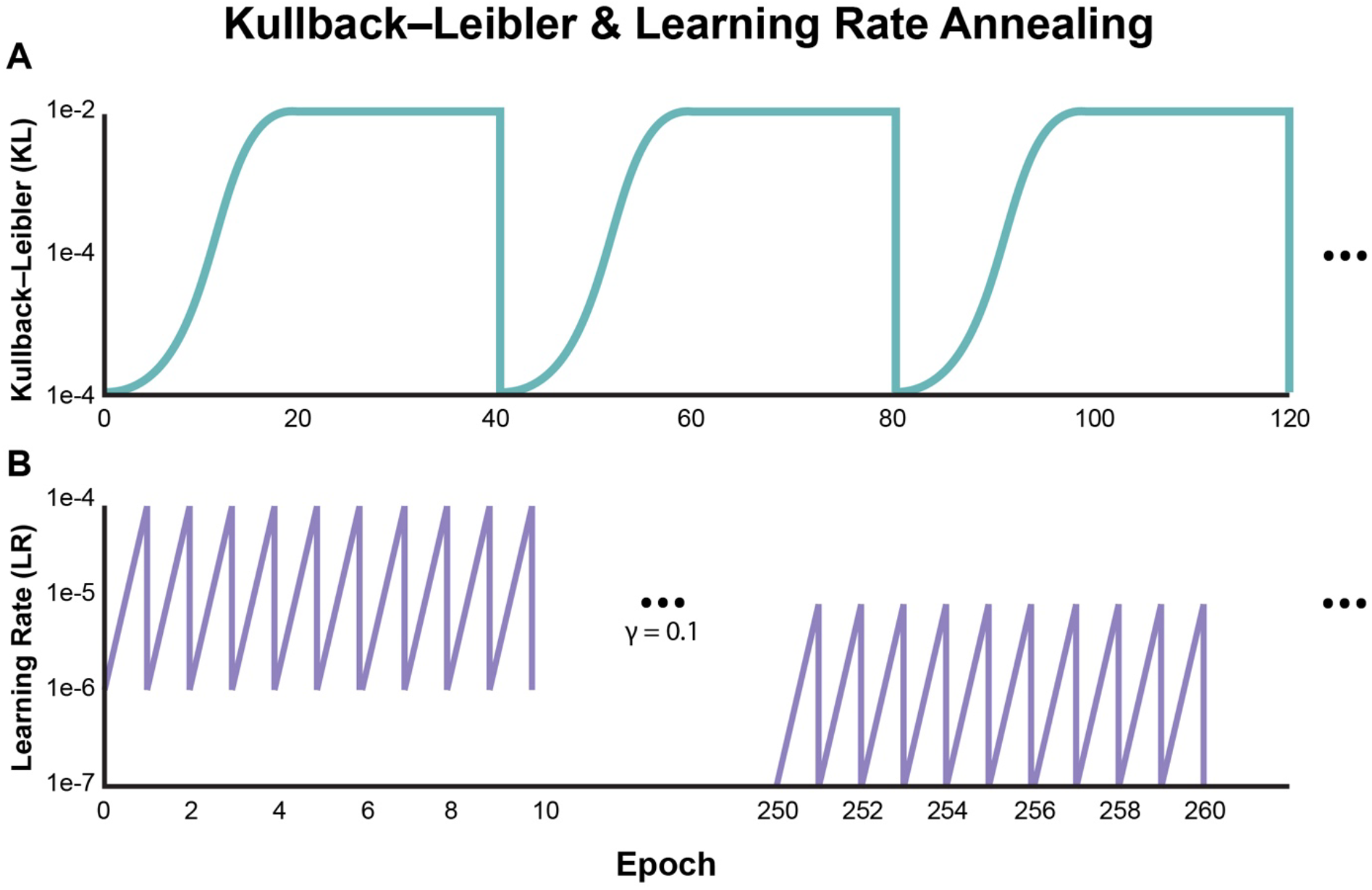
Kullback-Leibler (KL) and Learning Rate (LR) annealing schedules.

#### Brain-state interpretation: Dimensionality reduction & clustering

After model training has completed, the entire training and validation datasets (70/30%) are sequentially run through the model to get a continuous 512 dimensional latent-space representation of the data. These high-dimensional timeseries data are impossible for a human to interpret; thus, we require two steps before biologically meaningful analyses can be conducted: 1) Dimensionality reduction, 2) Brain-state clustering. To begin, dimensionality reduction is critical to interpreting the latent space architecture, but traditional methods like Principal Components Analysis (PCA) fail to separate brain-states into any discernible structure (**Figure 6A-B**). Thus we used Pairwise-Controlled Manifold Approximation and Projection (PaCMAP),^23^ a dimensionality reduction algorithm and successor to the popular Uniform Manifold Approximation and Projection (UMAP),^24^ that has demonstrated enhanced ability to capture global data structure, which is critical for analyzing large-timescale brain states (**Figure 6D**). Before feeding the latent space data into PaCMAP, we smoothed the 512-dimensional latent data with a 10-second averaging window, and 1 second stride - we found that this greatly increases the signal-to-noise ratio for the manifold approximation. The PaCMAP hyperparameters that differed from default were as follows: Distance: cosine, MN_ratio 2.0, FP_ratio 0.5 to enhance global structure definition of the reduced space.

**Figure 6:**
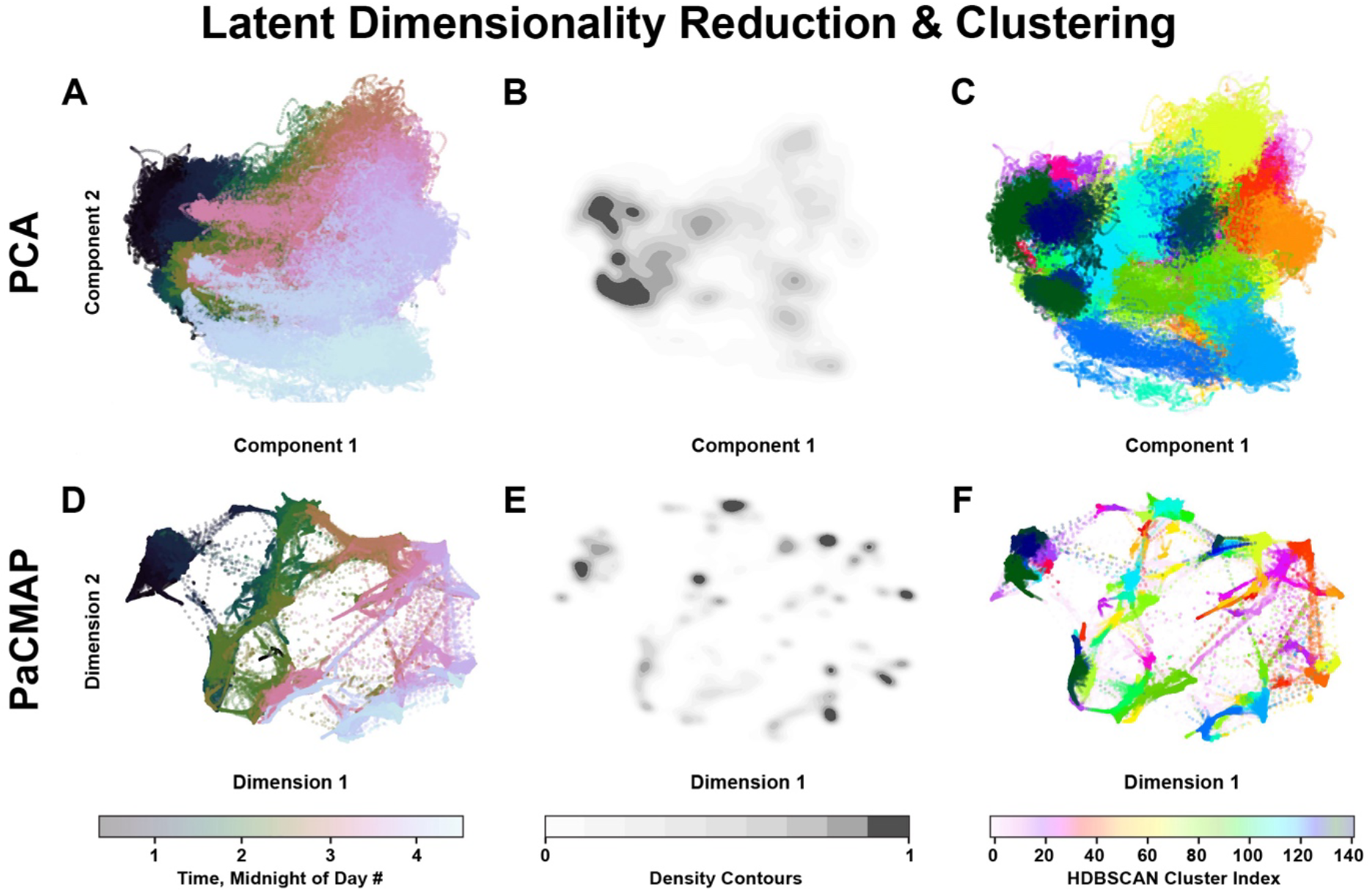
Latent dimensionality reduction and spatial clustering.

Next, differing densities of data appeared to be present in the data, as can be appreciated in the iso-density contours in **Figures 6B and 6E**. Thus, we clustered the PaCMAP reduced dimensional space using Hierarchical Density-Based Spatial Clustering of Applications with Noise (HDBSCAN).^25,26^ Of note, we conducted two separate runs of PaCMAP, the first is the 512 dimensional latent space reduced to a ten-dimensional space that was fed into HDBSCAN, and the second is a two-dimensional space for direct visualization in the plots shown. This was done to allow HDBSCAN to have access to an enriched ten-dimensional feature set that could help distinguished unique brain states without being limited to the simplified two-dimensional space. The hyperparameters that differed from default for HDBSCAN were as follows: min_cluster_size 200 (each data point is a 10 second average of latent data), min_samples 100. After dimensionality reduction and clustering, distinct brain-state groupings within the latent space become apparent in the 2D PaCMAP visualization (**Figure 6F**), that would have otherwise been indistinguishable by PCA (**Figure 6C**). Of note, the PaCMAP and HDBSCAN models were built with *only training data*.

Next, using the trained AR-βVAE, PaCMAP and HDBSCAN models, the completely withheld validation data can be run through the pipeline to observe the most-likely brain-state cluster for data the models have never previously seen. This flow of data through the pretrained models is how a real-time feedback signal would be generated for a closed-loop neuromodulation device. An ‘adaptive’ paradigm would arise from continuously updating these models based on new data to refine the brain-state estimation in real time.

### Brain-state model validation: perturbability & biological relevance

The generation of clustered latent spaces from epiphenomenal physiological timeseries has little meaning to potential closed-loop neuromodulation applications without two important forms of validation: 1) Evidence of brain-state perturbability with neurostimulation, 2) Evidence of biological relevance of brain-states. Both of these validation conditions must be reasonably met to proceed with a closed-loop neuromodulation paradigm. However, the exact criteria that dictates adequate validation is nebulous – especially establishing biological relevance because this is subject and disease specific. For example, the latent space of a person with epilepsy may differ greatly from a person with major depressive disorder. Or more nuanced, the difference between two people with epilepsy who experience different seizure types, take different medications, and are different ages. Thus, the following quantitative and qualitative techniques were implemented to capture the subject-to-subject variability in brain-state organization.

#### Brain-state perturbability with low-frequency neurostimulation

To capture possible effects of neurostimulation, we utilized our previously collected single-pulse electrical stimulation (SPES) paradigm conducted with 71 of our 118 pre-surgical epilepsy patients.^27^ This stimulation paradigm was not designed to maximize any possible brain-state modulation and consists of only 1 Hz, 300 microsecond biphasic pulses in trains of 10 seconds and at current levels of 1-5 milliamps. Thus, these SPES sessions offer a conservative insight into the effects of a low-energy stimulation paradigm’s effects on brain-state. The SPES data epoch (approximately 1-2 hours per subject) was excluded from training. Inference of the SPES epoch was conducted without manual stimulation artifact removal to minimize potential bias and simulate a real time device implementation – artifact is greatly mitigated by the ZHE normalization scheme and with the brevity of SPES artifact accounting for approximately 0.1-0.2% of any smoothed 10 second window. For continuous stimulation paradigms with a higher duty cycle, this technique may need to be amended.

To quantify any effects of neurostimulation, we implemented two high-level metrics: 1) number of brain-state transitions within a 60 second window, 2) number of unique brain-states occupied within a 60 second window. These metrics serve as a baseline validation to capture coarse information about the effects of neurostimulation. Importantly, due to the architecture of each model in a subject-specific manner, it is ill-posed to compare these metrics across subjects. Thus, we implemented a bootstrapping technique to estimate the statistical significance for each subject. Specifically, we evaluated the ‘state-transitions’ and ‘unique-clusters’ metrics for all non-overlapping 60-second windows within the SPES epoch, then calculated the metrics for an equivalent number of random 60-second epochs in the pre-SPES epoch. From these bootstrapped runs, 95% confidence intervals and two-sample t-tests can be conducted with Bonferroni-Holm multiple comparison correction.

#### Biological relevance of brain-states

Thus far, we have purposely left all methodology agnostic to any specific neurological disease. However, for the purpose of determining the biological relevance of the elucidated brain-states, we must interpret the latent spaces in the specific context of our cohort’s neurophysiological pathology – drug-resistant epilepsy. This type of validation relies on qualification of the latent space structure with detailed clinical information known about each subject. Most importantly for this cohort, the exact timing and type of all electroclinical seizures. With these events as ground truth, we can work backwards to overlay meaning to the brain-states that precede and follow these events.

With the epilepsy brain-state postulates in mind from **Table 1**, we asked three qualitative questions on the training data and validation data withheld from each subject’s trained model: 1) Does pre-ictal, ictal, and post-ictal activity in training data aggregate together in the latent space in a meaningful structure, and are any patterns based on different seizure types present? 2) Does completely withheld validation data for similar electroclinical epileptic events map to the expected latent space regions based on training data latent space assessment? 3) Can seizure propensity be assigned to the HDBSCAN clusters and plotted on a timeline for training data which can then be mapped to validation data?

To address the first and second validation questions, the entire training and validation datasets were run through the trained AR-βVAE model and the latent spaces for each subject were labeled with known ictal events. The latent space density iso-contours were concurrently plotted to assess the most common brain-states during the training and validation epochs. Next, the latent architectures during ictal and interictal periods were analyzed for meaningful aggregation of known electroclinical states. Finally, to assess the seizure propensity of the latent space clusters (i.e. brain-states), a timeline of cluster assignments for the training dataset was plotted against known ictal events. An exponentially decaying seizure propensity scoring mask (**Eq. 1**, where ‘t’ = sample count to seizure at sampling rate of 512 Hz, and d = decay factor of 20,000) was assigned to the timeline for all ictal events, with the maximum mask value at each timepoint was used to account for temporally adjacent ictal events.

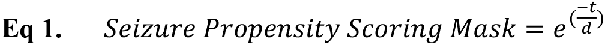

The brain-state clusters were then ranked by the mean scoring mask value of all time periods during which the cluster was assigned. With the cluster seizure propensity score assigned from the training dataset, the validation scored timeline can be plotted to assess the ability of the model to discern potentially high seizure risk brain-states on data the model has never seen before.

## RESULTS & DISCUSSION

The main goal of this text is to describe the proposed methods of brain-state modeling and provide an initial validation using high-level quantitative and qualitative metrics to assess proof-of-concept suitability for an adaptive closed-loop neuromodulation paradigm. Utilizing this modeling process, example latent spaces and brain-state clusters can be seen in **Figure 7**.

**Figure 7:**
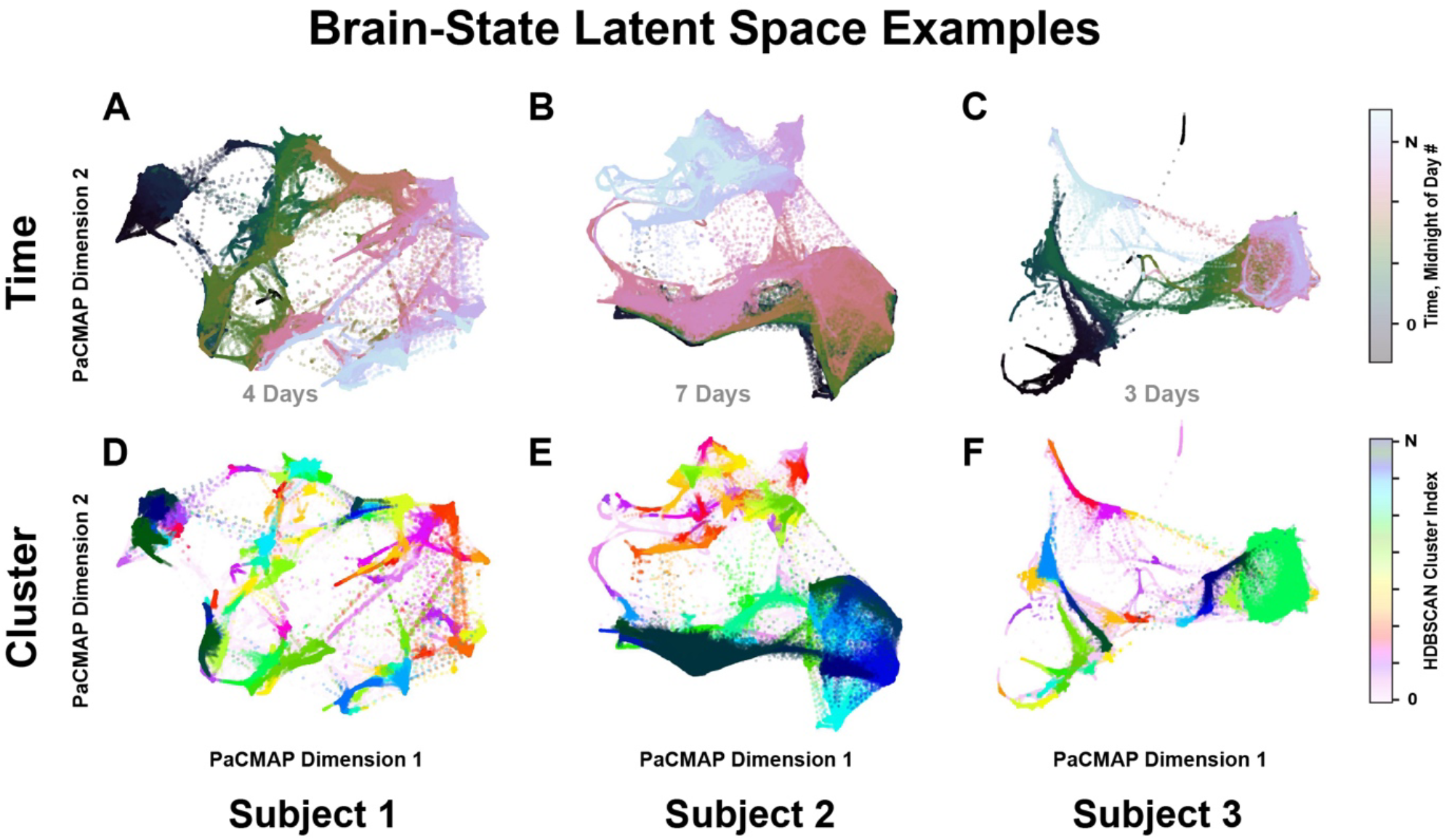
Example latent spaces depicting clustered functional brain-states.

### Brain-state observed to be significantly perturbed by low-frequency stimulation

The results of the neurostimulation perturbation analysis for these example subjects can be seen in **Figure 8**. SPES displayed evidence of significant brain-state perturbation, with the mean number of brain-state transitions observed to be 177-250% higher during the SPES epoch: Bootstrapping t-test p-values ranged from 6.35e-5 to 0.0367 for the average number of brain-state transitions during a given 60 second window in the pre-SPES vs. SPES epochs. Furthermore, the increase in brain-state transitions during the SPES epochs were not simply observed to be oscillations between the same brain-states because the number of unique brain-states increased by 127-163% during the SPES epochs, with p-values ranging from 6.74e-8 to 8.81e-3. These results indicate that even though SPES is a low-energy stimulation paradigm, it may still be significantly altering the brain-state as captured by the proposed brain-state modeling process.

**Figure 8:**
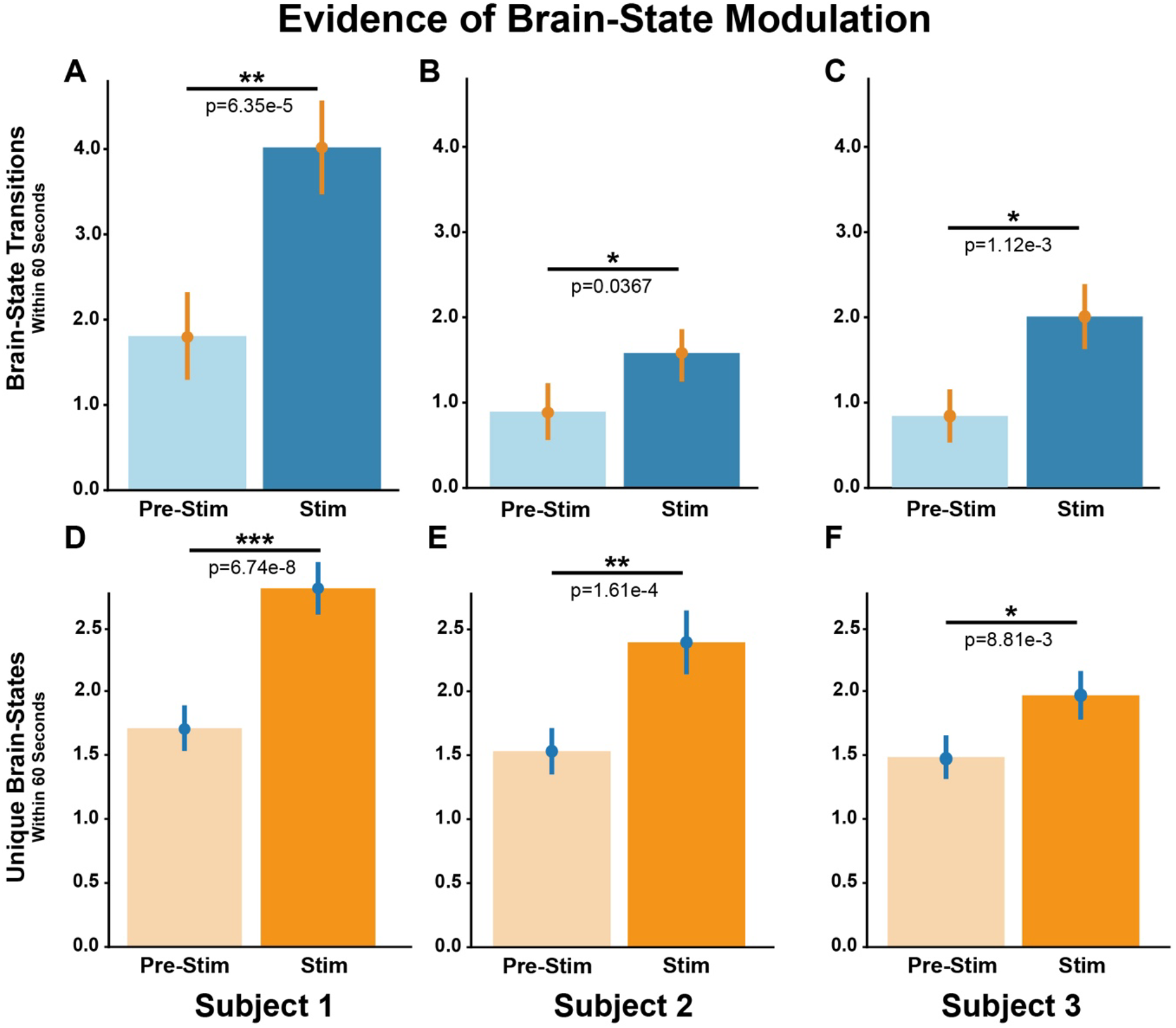
Significant perturbation of brain-states by single-pulse electrical stimulation.

### Brain-states appear to be biologically relevant based on inspection of latent space architecture

Beyond the ability to capture possible effects of neuromodulation, the brain-state modeling process must capture biologically meaningful information for a given neuropathology to be of clinical utility. To assess biological relevance, we have outlined the qualitative assessment of the latent spaces for three diverse subjects with epilepsy (**Figures 9-11**). Two subjects experienced focal to bilateral tonic-clonic (FBTC) seizures during SEEG recording (Subjects 1-2) and the third (Subject 3) experienced only subclinical and focal aware seizures (FAS). Of further interest, Subjects 1-2 had multiple seizure-free days prior to the first seizure captured on SEEG recording, whereas Subject 3 had multiple subclinical and FAS immediately on the first day of recording. These are important factors to consider when looking at the latent spaces for these subjects.

**Figure 9:**
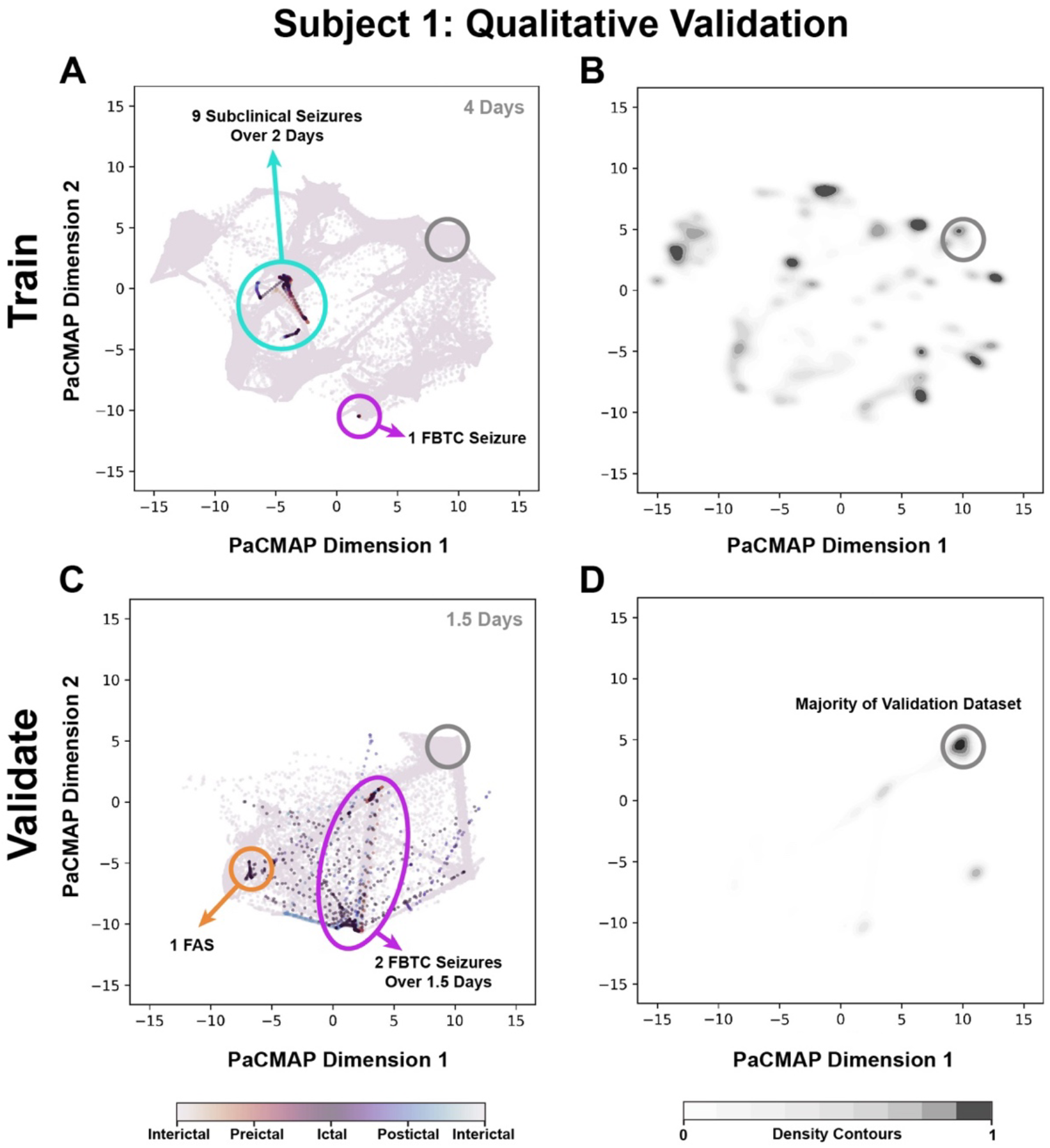
Training and validation latent spaces for Subject 1. Multiple seizure-free days prior to first ictal event. Only one FBTC seizure present in training dataset.

To begin, Subject 1’s latent space for the training data (**Figure 9A-B**) displays a distinct grouping of nine subclinical seizures that occurred over a two-day period. Furthermore, the sole FBTC seizure present in the training dataset is well separated from this subclinical aggregation, providing evidence that the preictal brain-state prodrome for the FBTC seizure differs to that of the subclinical seizures. However, these observations are for the data that the model had seen during training, thus analysis of the validation data completely withheld from the model training is of more interest. The validation latent spaces for Subject 1 (**Figure 9C**) at first appear noisy, but when looking at the density iso-contours of brain-states (**Figure 9D**), we observe that the majority of the validation brain-states are distant to areas of the latent space that were labeled as ictal in the training latent space. Furthermore, the two FBTC seizures present in the validation data mapped well to the training FBTC latent space despite only a single FBTC seizure being present for training. Finally, a new seizure type was present in the validation data (FAS) that mapped to a distinct location in the latent space near the subclinical aggregation despite this seizure type never having been seen by the model for training. This subject serves as a nice example of the potential variety in pre-ictal brain-states for different types of seizures, as outlined in the implications of **Postulate 1**.

The next example subject also had many seizure-free days prior to the first electroclinical event, however this patient’s training data had five FBTC seizures present in the training data (i.e. first 70% of data), thus this subject’s model was exposed to many more brain-state evolutions toward a FBTC seizure (**Figure 10A-B**). This is likely reflected in the tight aggregation of FBTC seizures in the validation data (**Figure 10C**) to the expected highest density of FBTC seizures in the training data (**Figure 10A**). Notably, the FTBC seizure in the training latent space that is most distant from the other four FBTC seizures was the first electroclinical event that the patient experienced during recording – perhaps triggering the subject to enter the high seizure propensity brain-states that resulted in four FBTC seizures over the next 2 days. An observation that provides further evidence for the subject’s overall brain-state shift throughout recording is the very different latent space occupancy during training (**Figure 10B**) compared to validation (**Figure 10D**). This serves as a good example of potentially different pre-ictal brain-state evolutions that can occur, as outlined in **Postulate 2**. Finally, this patient also experienced a focal impaired-aware seizure (FIAS) that mapped adjacent to the FBTC seizures in the training latent space, but no FIAS were present in the validation data to assess generalizability.

**Figure 10:**
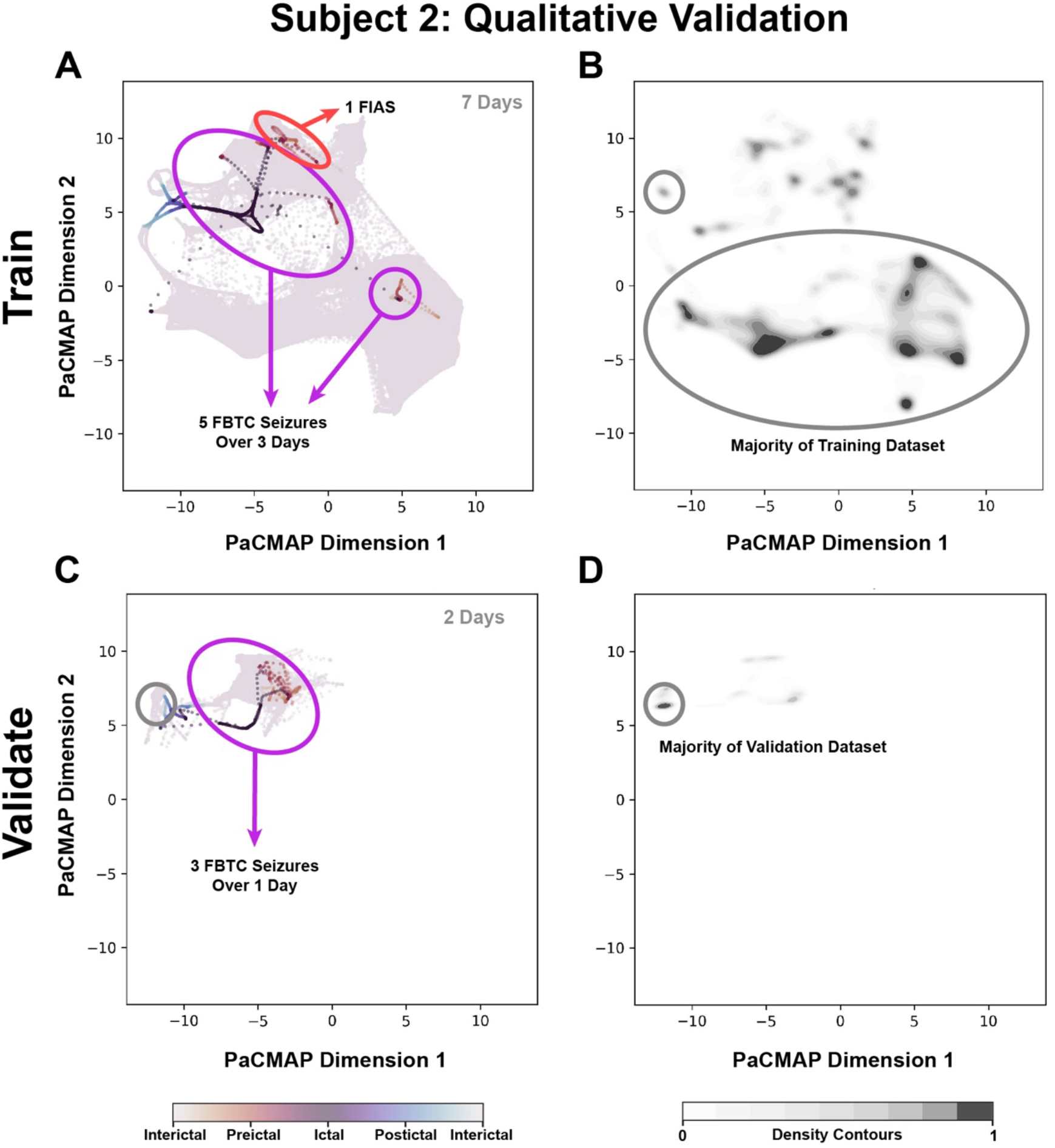
Training and validation latent spaces for Subject 2. Multiple seizure-free days prior to first ictal event. Multiple FBTC seizures present in training dataset.

The last example, Subject 3, had a significantly different electroclinical timeline of ictal events during SEEG recording, which is reflected in the training and validation latent spaces (**Figure 11**). The first observation is that the iso-density contours for training (**Figure 11B**) and validation (**Figure 11D**) are similar – this indicates that unlike Subjects 2 & 3, this subject likely did not experience a significant shift in the constellation of brain-states during recording. Thus, this subject serves as an exemplary case for the implications of **Postulate 3** that there may be no stable inter-ictal brain-state mapped into the latent space if the subject is experiencing a high ictal burden during recording. As suspected, unlike Subject’s 1 & 2 with a more concentrated aggregation of ictal activity in the latent space, Subject 3’s latent space is dominated by numerous FAS and subclinical seizures (**Figure 11A**). Importantly, the peri-ictal activity for the three subclinical and 15 FAS in the training data did map to different areas of the latent space, but the FAS tend to dominate the majority of the space. Evidence for this patient-specific hypothesis is observed in the validation latent space (**Figure 11D**), where the eight FAS present in the validation data occupy a similar, but also expansive, area of the latent space similar to the training data. For the purposes of brain-state modeling, this patient would have benefitted from a longer recording time to hopefully capture periods of lower seizure burden. Thus, as **Postulate 3** states, it may be difficult to extrapolate a ‘stable’ interictal brain-state that is far in latent space from ictal activity if no stable state has been previously observed.

**Figure 11:**
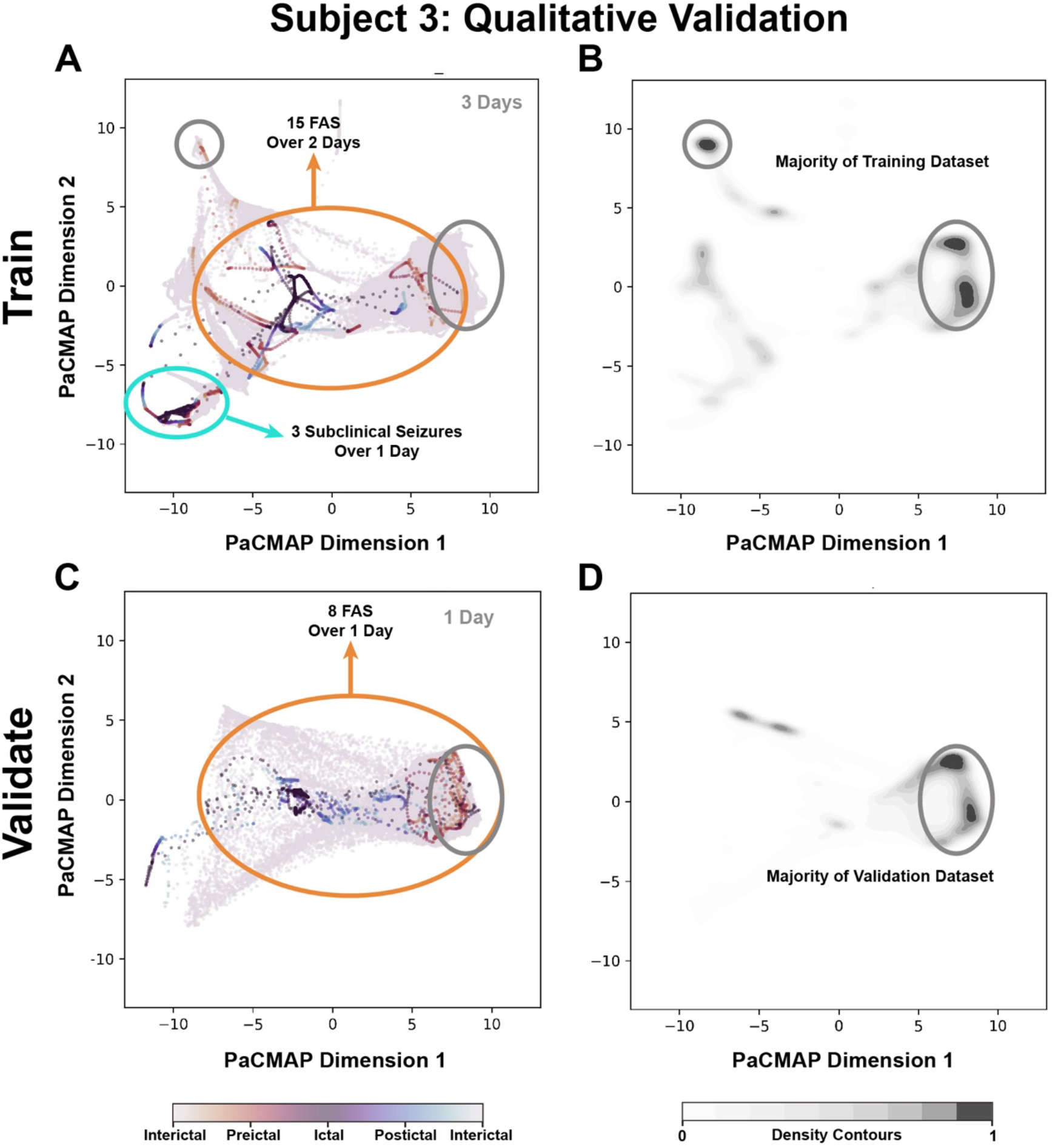
Training and validation latent spaces for Subject 3. Very high seizure burden during training and validation – likely no stable inter-ictal brain-states observed.

### Seizure propensity timeline demonstrates large-timescale transition to high ictal burden

An important feature of a potential closed-loop neuromodulation device for drug-resistant epilepsy is the ability to sense large-timescale shifts in brain-state. Without this ability, the device operates in one of two paradigms: 1) The “too little too late” paradigm where only an immediate pre-ictal signature can be detected and stim can be delivered, or 2) The device inappropriately characterizes large-timescale brain-states and thus operates in essentially an open-loop fashion with inadequate feedback signalling.

An example of the proposed brain-state modeling process ability to capture large-timescale brain-states is shown in **Figure 12** where example Subject 2’s seizure propensity timeline is plotted for training and validation data. This subject serves as a good example because the training data consists of many days of seizure-free recordings preceding multiple days of frequent FBTC seizures (**Figure 12A**), which were continued in the validation data (**Figure 12B**). Thus, using the seizure propensity masking technique (**Figure 12C**), the brain-state clusters can be ranked to align with estimated seizure-propensity within an arbitrary timeframe **(Figure 12E**), high values indicate high risk for a seizure. The validation timeline for this subject appropriately suggests that this patient spent most of the two days of validation recordings in a high seizure propensity state – which is in alignment with the two FBTC seizures this patient experience during validation. It is important to note that these are not immediately pre-ictal states and could be operated on by a neuromodulation device on the order of hours-days in an attempt to neuromodulate this subject through their mapped latent space to previously detected clusters with a lower seizure propensity.

**Figure 12:**
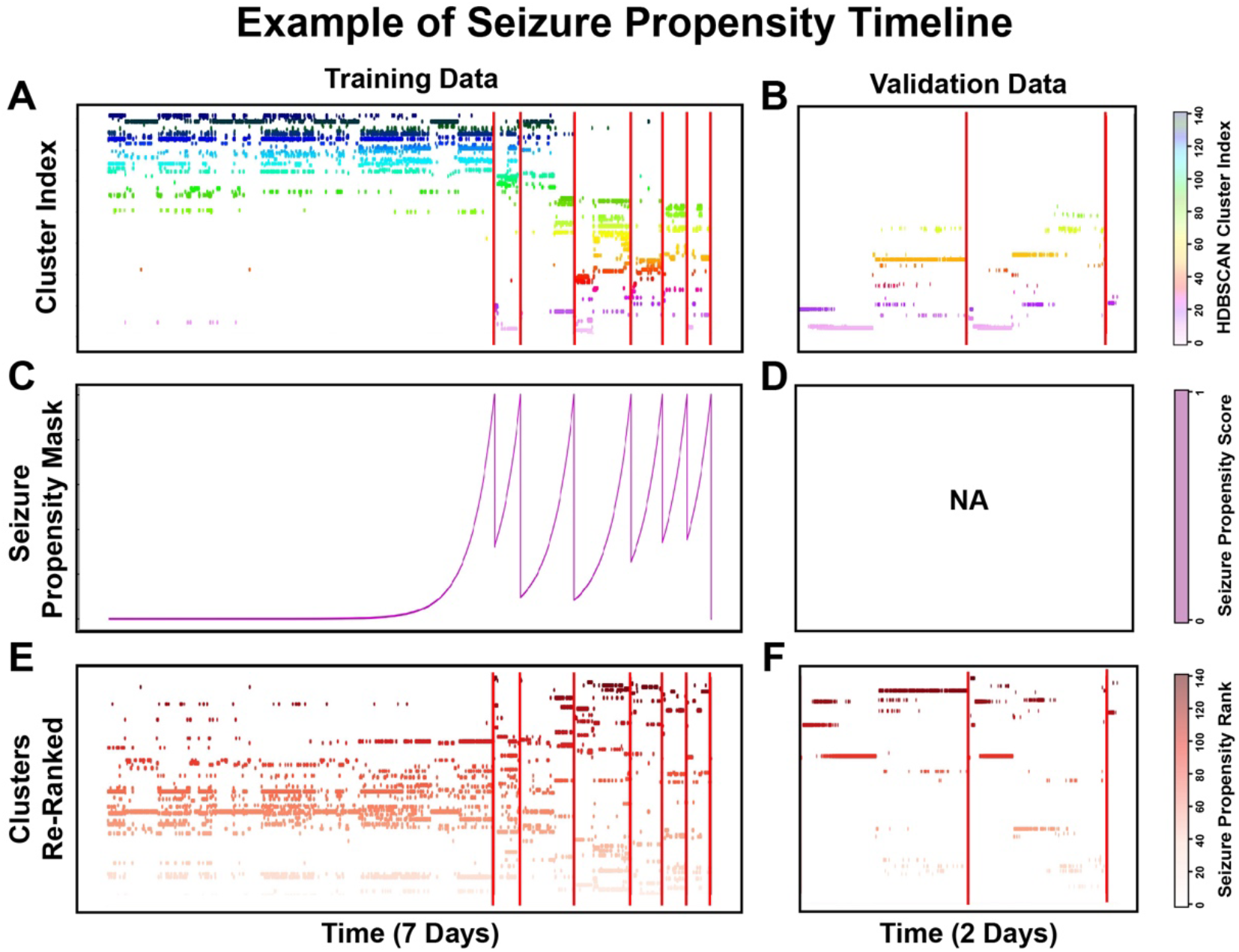
Example of Subject 2 seizure propensity timeline. Red vertical lines indicate ictal event.

